# Proteomic analysis of Stony Coral Tissue Loss Disease demonstrates coral-algal dysbiosis during disease progression

**DOI:** 10.64898/2026.09.01.748617

**Authors:** Shrinivas Nandi, Timothy G. Stephens, Kasey H. Walsh, Rebecca Garcia-Camps, Maria F. Villalpando, Rita I. Sellares-Blasco, Ainhoa L. Zubillaga, Haiyan Zheng, Aldo Croquer, Debashish Bhattacharya

**Affiliations:** Microbial Biology Graduate Program, Rutgers, The State University, New Brunswick, New Jersey, 08901, United States; Department of Biochemistry and Microbiology, Rutgers, The State University, New Brunswick, New Jersey, 08901, United States; Fundación Puntacana, Marine Innovation Centre, Puntacana Resort & Club, Punta Cana 23000, Dominican Republic; Fundación Dominicana de Estudios Marinos, Marine Research Dominican Republic, Calle Los Manantiales, Bayahibe, La Altagracia, 23000, Dominican Republic; EOHSI proteomics core, Rutgers, The State University, Piscataway, New Jersey, 08854, United States; The Nature Conservancy, Caribbean Division, Central Caribbean Program, Avenida 27 de Febrero, Plaza Central 1, Santo Domingo, 10148, Dominican Republic

**Keywords:** coral, *Diploria labyrinthiformis*, disease progression, multi-omics, proteomics, stony coral tissue loss disease

## Abstract

Stony Coral Tissue Loss Disease (SCTLD) has devastated Caribbean reefs, yet the host molecular response to infection is poorly understood. Previous gene expression studies of diseased corals identified shifts in the immune response, apoptosis, and coral-algal dysbiosis. Here, we characterized the proteomic response of *Diploria labyrinthiformis* to SCTLD by comparing protein abundance in healthy tissue from uninfected colonies and apparently healthy and neighboring diseased tissue from infected colonies. These results were compared with existing metagenomic data from the same samples that previously demonstrated significant shifts in the coral microbiome due to SCTLD. We identified 480 differentially abundant proteins when comparing diseased lesion and healthy tissues, but only 12 between apparently healthy and healthy tissues. Pathway-level analysis provides evidence of immune suppression in apparently healthy tissue, suggesting that host molecular responses precede visible disease progression. Diseased lesion tissues showed wound-healing responses combined with a decreased abundance of proteins involved in symbiosome maintenance and increased oxidative stress responses, consistent with host-algal dysbiosis. This result correlates with the previous metagenomic analysis of these samples which found that infected colonies exhibit distinct algal symbiont communities dominated by *Symbiodinium necroappetens*, whereas healthy colonies are dominated by *Durusdinium trenchii* and *Breviolum* spp. Comparison with existing transcriptomic studies revealed both shared and distinct molecular responses, underscoring the importance of integrating multi-omics approaches to understand coral diseases. Our results suggest that SCTLD in *D. labyrinthiformis* is associated with early immune suppression, coral-algal dysbiosis, oxidative stress, and subsequent wound-healing responses.

## Introduction

Over the past decade, coral reefs across the Caribbean have experienced a rise in disease outbreaks, with Stony Coral Tissue Loss Disease (SCTLD) emerging as one of the most severe and widespread (Alvarez-Filip et al., 2019; Brandt et al., 2021; Croquer et al. 2021 Papke et al., 2024). First reported in 2014 in Florida, SCTLD has since spread throughout the region, affecting over 22 coral species (Beavers et al., 2023; Precht et al., 2016). The disease is characterized by rapidly spreading lesions that often lead to colony death (Hawthorn et al., 2024). SCTLD is transmitted through direct contact and *via* the water column, which has contributed to its wide geographic spread, making it a major threat to Caribbean (and potentially beyond) reef ecosystems (Aeby et al., 2019; Muller et al., 2020; Rosenau et al., 2021; Studivan et al., 2022). Despite progress in characterizing SCTLD, key aspects of its etiology (Rosales et al., 2022, 2023), mechanisms of disease progression, and host immune responses (Beavers et al., 2023, 2025; Rossin et al., 2026) remain unresolved, which constrains the development of targeted interventions and effective mitigation strategies.

Early investigations of SCTLD largely focused on identifying bacterial pathogens using 16S rRNA amplicon sequencing, which revealed associations with several bacterial taxa (Arriaga-Piñón et al., 2024; Clark et al., 2021; Evans et al., 2023; Papke et al., 2024). Whereas antibiotic and probiotic treatments initially suggested a prokaryotic basis for the disease (Demko et al., 2025; Neely et al., 2020; Studivan et al., 2023; Ushijima et al., 2023), no single causative agent has been identified across all affected coral species (Heinz et al., 2024). Instead, current evidence points toward a more complex, polymicrobial etiology involving shifts in the coral microbiome (Huntley et al., 2022; Rosales et al., 2025). Increasingly, studies are expanding beyond bacteria to consider the broader holobiont, including viruses and microbial eukaryotes, which may play important roles in disease progression (Heinz et al., 2024; Nandi et al., 2025; Rosales et al., 2022; Veglia et al., 2022; Work et al., 2021). Observations of viral particles within algal endosymbionts (Work et al., 2021) and viral presence across different geographic sites and species have suggested that viral infections may play a prominent role in the onset of SCTLD (Nandi et al., 2025). Together, these findings highlight SCTLD as a multifactorial disease driven by dynamic interactions and opportunistic microbes within the coral microbiome (Meyer et al., 2019; Papke et al., 2024; Rosales et al., 2022).

The molecular mechanisms underlying SCTLD are still being actively investigated to better understand disease progression. Transcriptomic studies have elucidated the roles of coral immune response genes, extracellular matrix remodeling, wound healing and apoptosis pathways during infection (Beavers et al., 2023, 2025; Traylor-Knowles et al., 2021). These data suggest that SCTLD is associated with coral-algal dysbiosis. Transcriptomic profiling shows elevated expression of the RAB7 gene, a biomarker linked to disruption of the coral-algal symbiosis (Beavers et al., 2023; Traylor-Knowles et al., 2021). Elevated expression of peroxidase genes, an indicator of oxidative stress, may contribute to dysbiosis during SCTLD progression (Beavers et al., 2023; Papke et al., 2024; Traylor-Knowles et al., 2021). Histopathological studies support this hypothesis, demonstrating that coral-algal dysbiosis is associated with SCTLD-related tissue loss and lesion development (Rossin et al., 2026; Work et al., 2025).

Current analysis of SCTLD progression has relied heavily on transcriptomic data. However, studies of corals and other eukaryotes have shown a poor correlation between transcriptomic and proteomic data (Gygi et al., 1999; Williams et al., 2023). By measuring changes in translation products, proteomics provides a more direct assessment of the mechanisms underlying SCTLD infection responses. Protein abundance data capture regulatory processes beyond gene transcription, including post-transcriptional control and protein turnover, thereby establishing a mechanistic link between molecular regulation and phenotype that transcriptomics alone cannot address (Buccitelli & Selbach, 2020; Csárdi et al., 2015; Greenbaum et al., 2003). In this study, we extended our previous investigation of SCTLD-associated microbiome and virome dynamics in the Caribbean coral *Diploria labyrinthiformis* by examining host disease responses using proteomics. These data were used to identify the pathways impacted by an ongoing SCTLD infection.

## Methods

### Sampling plan and protein extraction

Tissue was collected from 10 *D. labyrinthiformis* colonies in waters near Bayahibe, Dominican Republic: five were healthy [**Figure 1A**] and 5 were SCTLD-infected colonies (see (Nandi et al., 2025) for details). In brief, from the five SCLD-affected colonies, samples were collected from the lesion site (hereinafter DL tissue) and ∼ 2 inches away from the diseased lesion in a region of the colony that appeared to be apparently healthy (hereinafter

**Figure 1:**
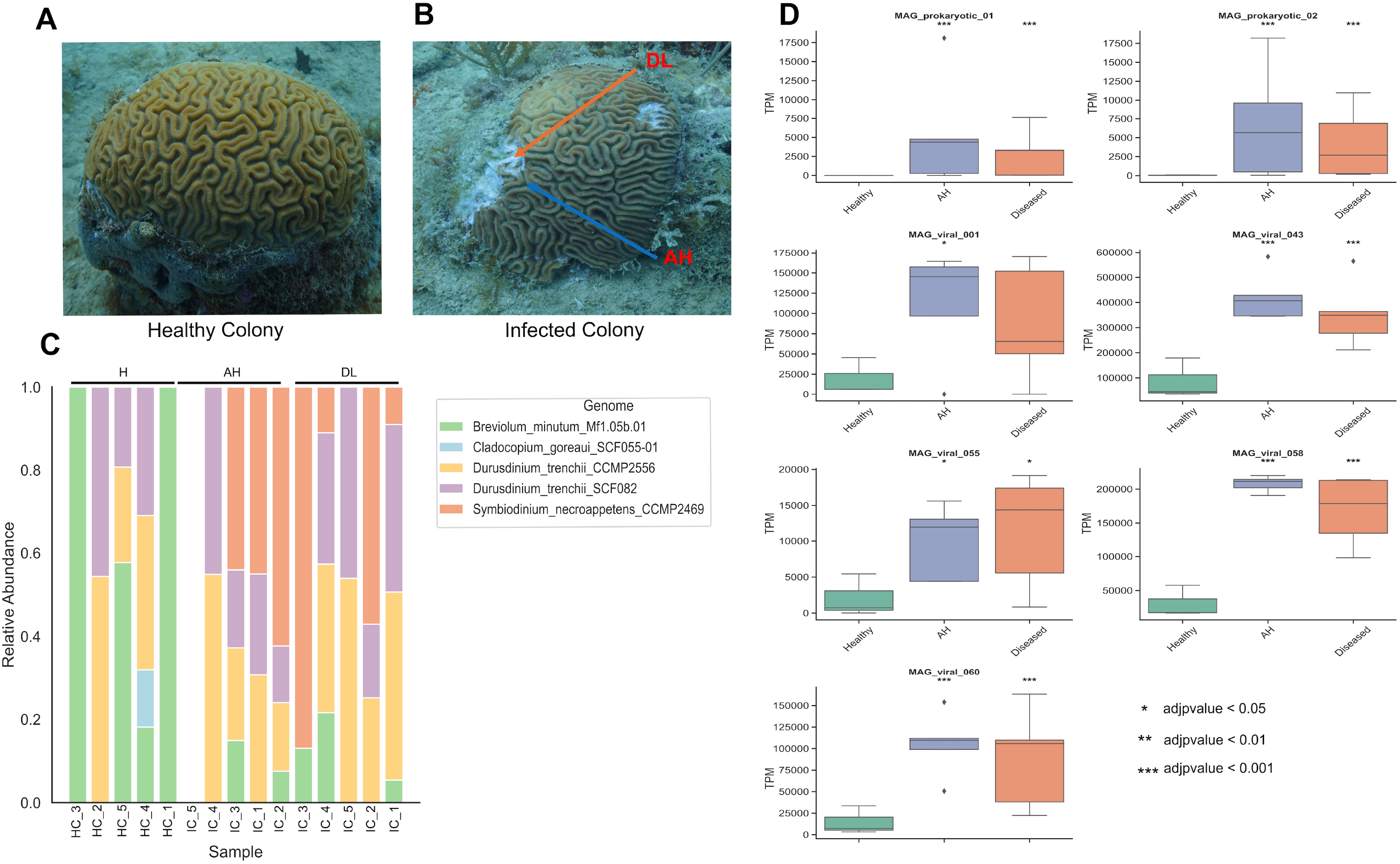
Summary of metagenomic analysis adapted from Nandi et al., (2025). **A)** Image of a healthy *Diploria labyrinthiformis* colony with no signs of SCTLD lesions. **B)** Image of a SCTLD-infected *D. labyrinthiformis* colony. The lower arrow indicates where the AH tissue sample was collected and the upper arrow shows where the DL tissue samples were collected. **C)** Stacked bar chart showing the relative abundance (Y-axis) of algal endosymbionts associated with sampled colonies (X-axis) based on mapped reads to genomes. One sample (IC_5) had no reads mapped to any of the endosymbiont genomes. IC: infected colony; HC: healthy colony. The bar above the plot indicates the tissue type for each sample. **D)** Box plots showing the TPM (Y-axis) of the SCTLD-associated MAGs in different tissue samples (X-axis). Stars on the top of each box plot show the statistical significance generated by DESeq2, compared to the healthy condition. A legend is presented on the bottom right of the panel.

AH tissue) [**Figure 1B**]. Collections from the five healthy colonies are hereinafter referred to as HC tissue. Samples were collected in triplicate (3 tissues x 5 colonies x 3 replicates = 45 samples). One sample was lost during transport (final samples n=44). After collection, samples were immediately stored in RNAlater and transported to Rutgers University, where they were stored at -80°C until processing.

Proteins were extracted using a RIPA lysis buffer, composed of Tris-HCl (50mM), NaCl (150mM), SDS (1%), and DI water. The buffer was chilled on ice until ready for use and one complete mini EDTA-free tablet was dissolved in 10 ml of lysis buffer immediately before extraction. A total of 500μl of ice-cold RIPA lysis buffer was added to a tube filled with 0.5mm silica beads. The samples were vortexed for at least five minutes, or until the skeleton was stripped from the tissue. The samples were then incubated on ice for 30 minutes, before centrifuging at 10,000 rcf for 10 minutes, followed by extraction of the supernatant. Protein concentration was assessed using a Bicinchoninic Acid Assay (Pierce BCA Protein Assay Kit Cat# 23225). Extracted proteins were stored in a freezer at -80°C.

### Proteomic data generation

Prior to data generation, a comprehensive metaproteomic database was assembled using: (i) *D. labyrinthiformis* predicted proteins (Metz et al., 2026); (ii) endosymbiont proteins, which were selected based on read mapping rates to the most relevant available genomes (Nandi et al., 2025), which included *Symbiodinium necroappetens* (González-Pech et al., 2021), *Cladocopium goreaui* (Chen et al., 2022)*, Breviolum minutum* (Shoguchi et al., 2013), and *Durusdinium trenchii* (Dougan et al., 2024) [**Figure 1C**]; (iii) microbiome MAGs generated in (Nandi et al., 2025). LC-MS/MS analysis was performed at the Proteomics Core at Rutgers University. 10ug of sample were diluted into 100 ul lysis buffer (2% SDS, 50 mM HEPEs, pH 8, 50 mM EDTA) and subjected to reduction with 5 mM DTT for 30 min at 60C, alkylation with 20mM iodoacetamide for an hour at room temperature in the dark and followed by SP3 beads digestion method (Hughes et al., 2019) with trypsin (sequencing grade, Thermo Scientific Cat#90058) in 100mM ammonium bicarbonate, 2 mM CaCl2 and incubated at 37^0^C for overnight. Peptides acidified with formic acid and volume was reduced to 20 ul by vacuum, then 100 ul 50% acetonitrile, 0.1 % TFA were added to the samples and frozen at -80^0^C for at least 30 minutes. The samples were centrifuged at 25,000 g for 10 min. The supernatant was dried in vacuum concentrator until volume was less than 10 ul. The samples were desalted using stage tip method (Rappsilber et al., 2007) and 10% of the sample was analyzed by nanoLC-MSMS.

### Liquid chromatography-tandem mass spectrometry (LC-MS/MS)

Samples were analyzed by LC-MS using a Nano LC-MSMS (Dionex Ultimate 3000 RLSCnano System, Thermofisher) interfaced with Eclipse (Thermofisher). Samples were loaded onto a fused silica trap column Acclaim PepMap 100, 75umx2cm (ThermoFisher). After washing for 5min at 5µl/min with 0.1% TFA, the trap column was brought in-line with an analytical column (Nanoease MZ peptide BEH C18, 130A, 1.7um, 75umx250mm, Waters) for LC-MS/MS. Peptides were fractionated at 300nL/min using a segmented linear gradient 4-15% B in 30min (where A: 0.2% formic acid, and B: 0.16% formic acid, 80% acetonitrile), 15-25% B in 40min, 25-50% B in 44min, and 50-90% B in 11min. Solution B then returns at 4% for 5min for the next run.

Raw data were analyzed with the custom protein database as described previously (i.e., host, algal symbionts, and microbiome) for Direct DIA searching using Spectronaut v1.9 (Biognosys AG) with recommended settings. Protein Group quantity was calculated and normalized across the experiment using the MaxLFQ method. Proteins in each group were filtered based on a posterior error probability (PEP < 0.01) and a protein group *q*-value threshold (FDR < 0.01).

### Proteomic data analyses

Proteomic data analysis was performed in R (v4.5.1) using the *Protti* package (v0.9.1) (Quast et al., 2022). Protein Groups were first separated into host, algal, and microbiome subsets for independent processing. When a Protein Group contained multiple protein IDs, KEGG annotations were evaluated for each individual protein. Protein Groups were retained only if all constituent proteins were assigned the same KEGG Orthology (KO) annotation (hereinafter referred to as “proteins”). Potential contaminants provided by the LC-MS facility were removed. The data matrix was further filtered to exclude observations with intensity values below 5, which likely represented false assignments. Protein intensities were log₂-transformed using the *mutate* function from *dplyr* and base R’s *log2* function. Outlier samples were identified using PCA and data completeness assessments via the *qc_pca* and *qc_data_completeness* functions [**Supplemental Figure 1**]. Based on these analyses, eight samples (P115, P119, P121, P132, P133, P134, P136, and P138) were removed. Protein groups missing from more than 80% of samples were also removed, which is a commonly used threshold in proteomic and metabolomic studies (Chille et al., 2026; Nandi et al., 2026). Any missing values in the remaining proteins were imputed using the *missForest* function (package *missForest*) with default parameters (*maxiter* = 10, *ntree* = 100) and a random seed of 124. Log₂-transformed peak intensities were then median-normalized within each tissue group (HC, AH, or DL) using the *normalise* function in *protti*, yielding the final cleaned protein abundance matrix. Differential abundance was calculated using *protti*’s *calculate_diff_abundance* function, with HC tissue samples as the control group and DL or AH tissues as the treatment group. Statistical significance was determined using the *moderated_t_test*, with *p-values* adjusted for multiple testing using the Benjamini-Hochberg (BH) method. Proteins were considered differentially abundant (hereinafter differentially abundant proteins [DAPs]) if they had an |log₂ fold change| > 0.5 and adjusted *p*-*value* < 0.05.

KEGG annotations were assigned to proteins by GhostKOALA (v 3.1) (Kanehisa et al., 2016). DAPs that had no KEGG annotation were subsequently compared against the nr database using DIAMOND blastp (*e*-value 1e-05) (Buchfink et al., 2015). DAPs were grouped into KEGG pathways at the C-Description level, with enrichment of these pathways assessed using Gene Set Enrichment Analysis (GSEApy v1.0.6) (Fang et al., 2023). The *t-statistic* was utilized from the *protti* differential abundance analyses to pre-rank the proteins. The analyses were performed using gp.prerank (perumation_num 1000, seed 42, min_size 10, and max_size 1000). A pathway was considered enriched if it had a Normalized Enrichment Score (NES) > 1.0 and FDR *q*-value < 0.05. A pathway was considered depleted if it had a NES < -1.0 and FDR *q*-value < 0.05. All downstream analyses were performed in Python v3.9.7 using the base packages, Pandas (v2.2.3), and NumPy (v1.26.4). Plots were generated using Matplotlib (v3.6.2) and Seaborn (v0.12.1).

## Results

### Proteomic data clean up and normalization

Proteins were initially partitioned by biological origin into host, endosymbiont, and microbiome fractions. Endosymbiont-derived proteins were excluded from downstream analysis for the following reasons: (i) endosymbiont cell-count data were unavailable, preventing correction of proteomic shifts caused by changes in endosymbiont biomass between samples (Nandi et al., 2026), and (ii) based on genomic read mapping rates, endosymbiont communities differed sharply between healthy and SCTLD-affected colonies, with infected colonies dominated by the opportunistic *Symbiodinium necroappetens*, whereas healthy colonies were dominated by *Durusdinium trenchii* and *Breviolum* spp. [**Figure 1C**; (Nandi et al., 2025)]. These factors confound interpretation of endosymbiont protein abundances and necessitated their removal from downstream analyses.

Prior to quality control, 8,946 host-derived proteins were identified in the *D. labyrinthiformis* data. Host proteins were filtered for high-confidence identifications using a PEP score < 0.01 and PG Q value < 0.01, resulting in the retention of 7,713 proteins. Proteome completeness was evaluated per sample; four samples (P121, P115, P119, and P136) that exhibited <70% completeness were removed **[Supplemental Figure 1A]**. To identify additional outliers, hierarchical clustering [**Supplemental Figure 1B**] and principal component analysis (PCA) were performed [**Supplemental Figure 1C**]. Both approaches flagged an additional four samples (P132, P133, P134, and 138) which clustered with the low completeness samples and were thus removed from downstream analysis. This resulted in a final tally of 36 samples after cleaning, removing four AH samples, one DL sample and three healthy samples.

Following filtering for missingness, 7,146 proteins were retained and had missing values imputed. Quality control diagnostics confirmed improved sample completeness and clustering structures after outlier removal and missing value imputation **[Supplemental Table 1**].

The same data-cleaning and filtering pipeline described above was applied to the microbiome-derived proteome. Overall, the microbiome used for this analysis had 301,571 proteins, however only 2,240 proteins were detected in the proteomic data. Furthermore, overall data completeness was exceptionally low, with only three samples exhibiting completeness > 50% [**Supplemental Figure 2**]. The combination of poor sample completeness and high protein attrition after QC, rendered the dataset unsuitable for analysis to make robust biological interpretations. As a result, subsequent analyses focused exclusively on the coral host proteome, integrating findings from the prior metagenomic results, to assess the molecular response associated with active SCTLD infection.

### DAP analysis

We identified 480 differentially abundant proteins (DAPs) which were classified as increased in abundance if they had a log_2_FC > 0.5 and an adjusted *p*-value < 0.05 and decreased in abundance if they had a log_2_FC < −0.5 and an adjusted *p*-value < 0.05 (Xie et al., 2026). Of the 7,146 proteins that remained after filtering, 12 DAPs distinguished the AH and HC samples, of which two had an increased abundance and 10 had a decreased abundance in the AH samples when compared to HC [**Figure 2A, Supplemental Table 2]**. In the comparison between DL and HC samples, 480 DAPs were identified, of which 155 had increased abundance and 325 had decreased abundance [**Figure 2B**]. DAPs appeared to show similar trends in both AH and DL, suggesting progressive protein abundance changes [**Figure 2C, Supplemental Table 2**].

**Figure 2:**
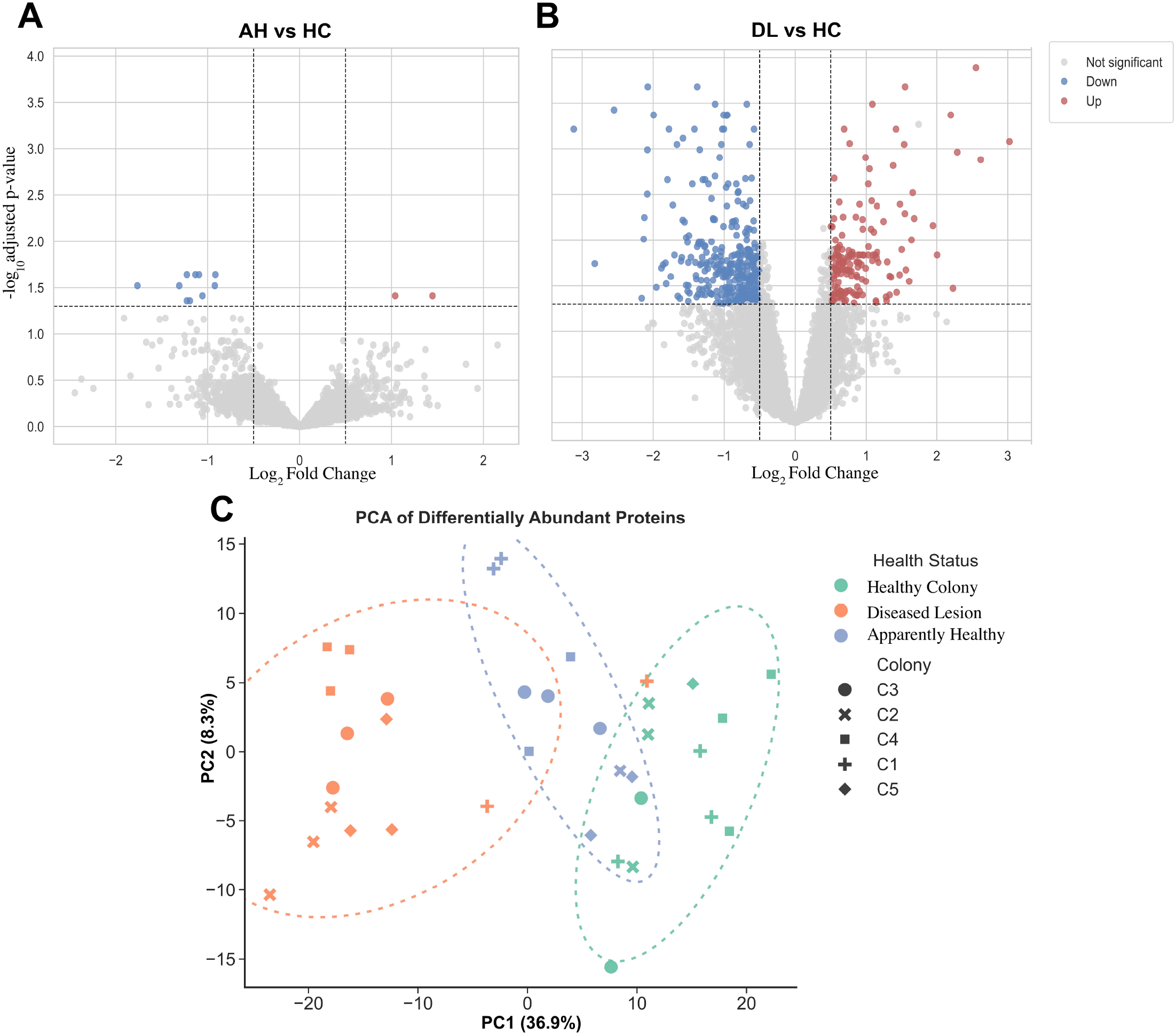
Differentially abundant proteins in diseased, relative to healthy colony tissue. **A)** Volcano plot showing the DAPs between AH vs. HC. Each point represents a protein. The X-axis shows the Log_2_ fold change and the Y-axis shows the -log_10_(adjusted *p-value*). Bold lines show the cut offs utilized i.e., |Log_2_FC| > 0.5 and adjusted *p-value* < 0.05. The red points show proteins with increased abundance in AH compared to HC, blue points show proteins with decreased abundance in AH compared to HC, and the grey points are the non-DAPs. **B)** Volcano plot showing the DAPs between DL and HC; features are same as in panel **A. C)** Principal component analysis of normalized abundances of differentially abundant proteins. Each point represents an individual tissue sample. Samples are coloured by health status and shaped by colony identity. Dotted ellipses represent two-standard-deviation covariance ellipses calculated separately for each health-status group. The percentages of total variance explained by PC1 and PC2 are shown on the X- and Y-axes, respectively. PCA of all proteins is presented as Supplemental Figure 1C.

A total of 480 DAPs were identified across all comparisons (AH vs. HC and DL vs. HC), 264 of which had assigned KEGG annotations. Of the remaining 216 which did not have assigned KEGG annotations, 107 returned informative BLAST hits against the nr database, the remaining 136 proteins returned non-informative descriptions, including uncharacterized (n = 99), hypothetical (n = 7) or did not return hits to the database (n = 3) **[Supplemental Table 2**]. Because the latter proteins did not have assigned functional annotations, they were excluded from downstream analyses. Nonetheless, these findings highlight the potentially important roles of functionally uncharacterized “dark” proteins in the coral response to SCTLD (Stephens et al., 2026).

Only 11 DAPs were identified in the AH vs HC comparison. Notably, these proteins also showed a significant response in the DL samples. Seven of these proteins had descriptive annotations assigned through comparison against nr, of which, two were copies of the 1-failed axon connections homolog protein, which had a mixed response, with one copy increasing in abundance (FC = 1.44, adj-*p-value* = 0.038) and the other decreasing (FC = - 1.09, adj-*p-value* = 0.022). A complex III assembly factor LYRM7 protein decreased in abundance (FC = -1.13, adj-*p-value* = 0.022), and the immune response associated gene perlucin-like protein also decreased in abundance (FC = -1.17, adj-*p-value* = 0.030). Lastly, a dentin sialophosphoprotein-like protein associated with both immune response and skeletal structure (Goldberg et al., 2011; Levy & Mass, 2022) also decreased in abundance (FC = - 1.22, adj-*p-value* = 0.022).

### Extracellular matrix associated proteins

The DL vs. HC DAPs is summarized in **Figure 3A** [**Supplemental Table 2**]. Collagen-associated proteins, which play essential roles in coral wound healing (Beavers et al., 2023; Papke et al., 2024; Rahman, 2019; Traylor-Knowles et al., 2021), showed mixed responses. Specifically, four collagen-associated proteins decreased in abundance, whereas six increased. In addition, prolyl 4-hydroxylase alpha subunit (P4HA), a protein involved in collagen synthesis, also increased in abundance (K00472; FC = 0.89, adj-*p-value* = 0.017). Four copies of hemicentin (K17341) decreased in abundance, whereas three copies of cartilage intermediate layer protein 1 increased in abundance. The actin cytoskeleton-regulatory complex protein decreased in abundance (FC = -0.63, adj-*p-value* = 0.003), as did the cytoskeleton-associated protein MEMO1 (K06990; FC = -1.62, adj-*p-value* = 0.043).

**Figure 3:**
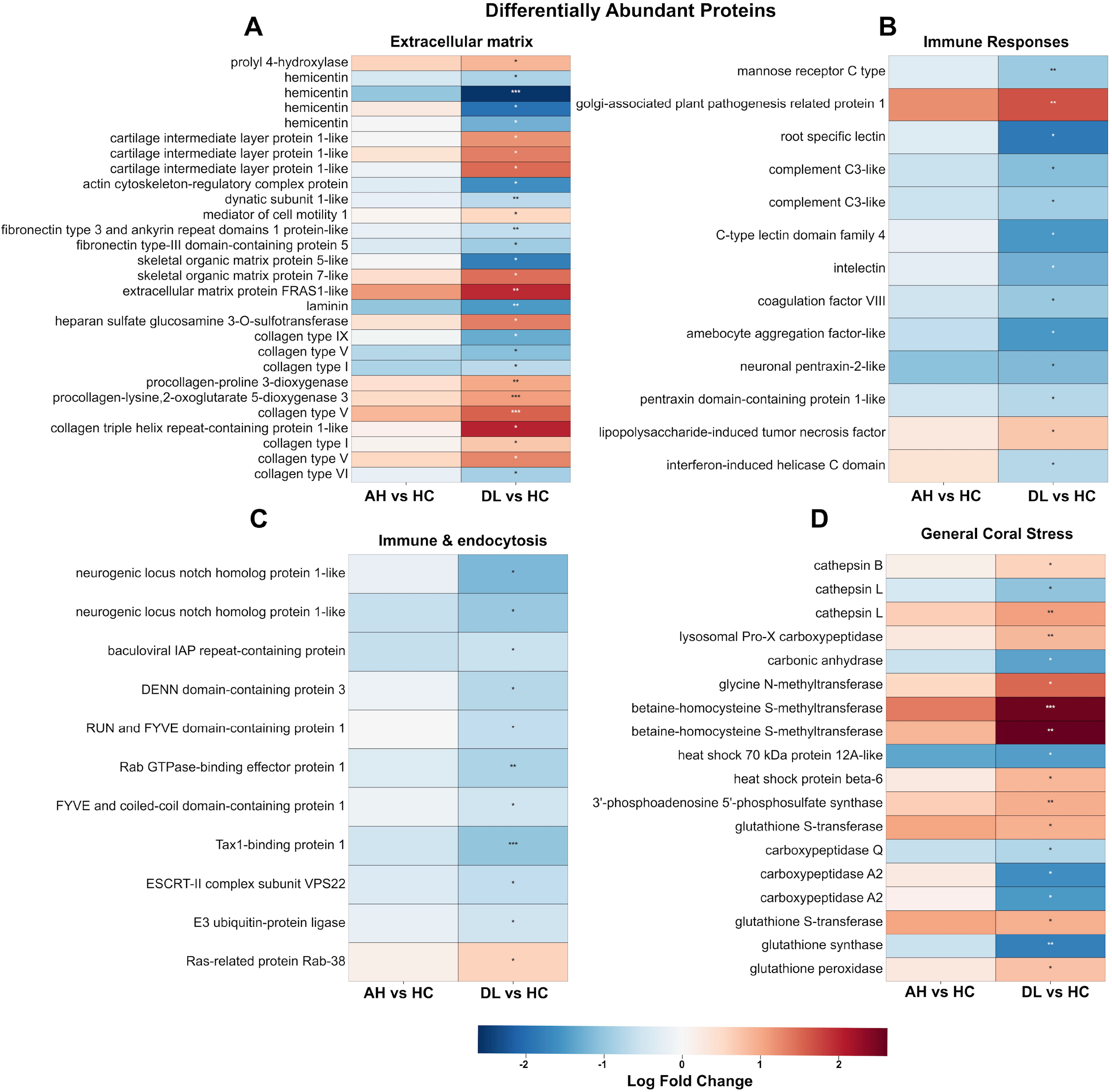
Differentially abundant proteins within functional categories previously associated with SCTLD. Four heatmap panels show the log_2_fold changes in protein abundance between AH vs. HC and DL vs. HC (X-axis). Proteins are grouped into four functional categories: (**A)** extracellular matrix (ECM), (**B)** immune responses, **(C)** immune and endocytosis, and (**D)** generalized coral stress responses. The colour scale has been presented in the figure (red represents an increase in protein abundance in AH or DL treatments compared to HC, blue represents a decrease). Asterisks denote the statistically significant differential abundance after multiple testing correction (* adjusted *p-value <* 0.05, ** < 0.01 and *** < 0.001).

Several additional ECM-related proteins were differentially abundant. Fibronectin type 3 and ankyrin repeat domain-containing protein 1 showed decreased abundance (FC = -0.63, adj-*p-value* = 0.036), as did leucine-rich repeat and fibronectin type III domain-containing protein 5 (FC = -0.93, adj-*p-value* = 0.046), and a laminin protein (a core ECM protein; K05635; FC = -1.49, adj-*p-value* = 0.008). Two uncharacterized skeletal organic matrix proteins exhibited opposing responses, with one decreasing (FC = -1.81, adj-*p-value* = 0.029) and the other increasing in abundance (FC = 1.43, adj-*p-value* = 0.037). Conversely, the skeletal ECM protein FRAS1-like increased in abundance (FC = 1.94, adj-*p-value* = 0.006), as did a heparan sulfate-glucosamine 3-sulfotransferase 3 protein (K07809; FC = 1.33, adj-*p-value* = 0.024).

### Coral immune associated proteins

Coral immune responses to SCTLD have been extensively characterized (Beavers et al., 2023; Papke et al., 2024; Traylor-Knowles et al., 2021). The DAPs identified in the DL tissue primarily showed evidence of immune suppression, summarized in **Figure 3B**. We observed three different lectin proteins all of which showed a decreased abundance: root-specific lectin-like (FC = -1.85, adj-*p-value =* 0.018), C-type lectin domain family 4 (K10059: FC = - 1.51, adj-*p-value =* 0.032), and intelectin (K17527: FC = -1.25, adj-*p-value =* 0.033). Two copies of complement factor, C3 had a decreased abundance (K03990: FC = -1.11, adj-*p-value =* 0.023 and FC = -0.91, adj-*p-value =* 0.033). The coagulation factor VIII protein also showed a decreased abundance (K03899: FC = -0.98, adj-*p-value =* 0.033). We also observed a decreased abundance of mannose receptor type C (CD206) (FC = -0.94, adj-*p-value =* 0.002), which are associated closely with lectins (Guo et al., 2025).

We also observed a decreased abundance in a NOD-like receptor family protein NLRC3 (K22614, FC = -0.56, adj-*p-value =* 0.017), a hemagglutinin/amebocyte aggregation factor-like protein (FC = -1.50, adj-*p-value e =* 0.010), a IFIH1 protein (K12647: FC = -0.74, adj-*p-value* = 0.042), and two pentraxin domain containing proteins (neuronal pentraxin-2-like [FC = -1.15, adj-*p-value =* 0.027] and sushi, von Willebrand factor type A, EGF and pentraxin domain-containing protein 1-like [FC = -0.74, adj-*p-value =* 0.034]. An increased abundance was observed in a LITAF protein (K19363: FC = 0.73, adj-*p-value* = 0.0021).

Proteins associated with autophagy, apoptosis, and endo-lysosomal trafficking are implicated in SCTLD and stress responses, which are shown in **Figure 3C** (Beavers et al., 2023, 2025). We observed that two copies of the cell fate regulator NOTCH1 showed a decreased abundance (K02599: FC = -1.18, adj-*p-value* = 0.039 and FC = -1.00, adj-*p-value* = 0.021). We also observed a decreased abundance of baculoviral IAP repeat-containing protein (K16060: FC = -0.58, adj-*p-value* = 0.037). Several proteins associated with endosomal trafficking and autophagy showed a decreased abundances: DENN domain-containing protein 3 (K20162: FC = -0.75, adj-*p-value* = 0.042), Rabaptin-5 (K12480: FC = -0.81, adj-*p-value* = 0.004), RUFY1/RABIP4 (K12482: FC = -0.65, adj-*p-value* = 0.028), FYCO1 (K21954: FC = -0.53, adj-*p-value* = 0.046), TAX1BP1 (K21347: FC = -1.03, adj-*p-value* < 0.01), SNF8 (K12188: FC = -0.64, adj-*p-value* = 0.014), and leucine-rich repeat and sterile alpha motif-containing protein 1 (K10641: FC = -0.54, adj-*p-value* = 0.037). In contrast, RAB38, which is associated with lysosome-related organelle biogenesis, showed an increased abundance (K07923: FC = 0.60, adj-*p-value* = 0.013), as did the Golgi plant-pathogenesis related protein (FC = 1.66, adj-*p-value* = 0.003).

### Proteins associated with endosymbiosis and coral stress

We observed decreased abundance in proteins associated with the symbiosome and maintenance of the coral-algal symbiosis [**Figure 3D**] (Maruyama et al., 2026). We observed an increased abundance of cathepsin B (K01363: FC = 0.60, adj-*p-value* = 0.021). In contrast, we observed a mixed response for cathepsin L, wherein one copy had a decreased abundance (K01365: FC = -1.05, adj-*p-value* = 0.014) and the other had an increased abundance (FC = 1.07, adj-*p-value* = 0.0017). We observed four carboxypeptidases, of which three had a decreased abundance including, two copies of carboxypeptidase A2, CPA2 (K01298: FC = - 1.60, adj-*p-value* = 0.016 and FC = -1.50, adj-*p-value* = 0.031) and carboxypeptidase Q like, CPQ (K01302: FC = -0.79, adj-*p-value* = 0.0017). One copy of lysosomal pro-x carboxypeptidase, PRCP had an increased abundance (K01285: FC = 0.85, adj-*p-value* = < 0.001). Finally, we observed a decreased abundance of carbonic anhydrase (K01672: FC = - 1.41, adj-*p-value* = 0.017).

We also observed proteins previously associated with coral stress in our set of HC vs. DL DAPs. An increased abundance was observed for two copies of betaine-homocysteine methyltransferase (BHMT, K00544: FC = 2.55, adj-*p-value* = < 0.001 and FC = 2.61, adj-*p-value* < 0.001), one glycine N-methyltransferase protein (GNMT, K00552: FC = 1.50, adj-*p-value* = 0.012), and multiple reactive oxygen species (ROS) detoxification proteins, including, glutathione S-transferase (K00799: FC = 0.94, adj-*p-value* = 0.026) and glutathione peroxidase (K00432: FC = 0.77, adj-*p-value* = 0.014). Notably, glutathione synthetase had a decreased abundance (K21456: FC = -1.79, adj-*p-value* = 0.002). Lastly, two heat-shock associated proteins showed mixed responses, notably HSP70 had a decreased abundance (FC = -1.46, adj-*p-value* = 0.015) and HSP6 had an increased abundance (FC = 0.87, adj-*p-value* = 0.039).

### Pathway enrichment analyses

Gene set enrichment analysis was performed using KEGG pathway annotations at the C-description level. Of these, 10 pathways were enriched and 10 pathways were depleted in AH when compared to HC [**Figure 4A; Supplemental Table 3**]. Notably the carbohydrate digestion and absorption pathway was enriched (ko04973: NES = 1.90, *q*-value = 0.027).

**Figure 4:**
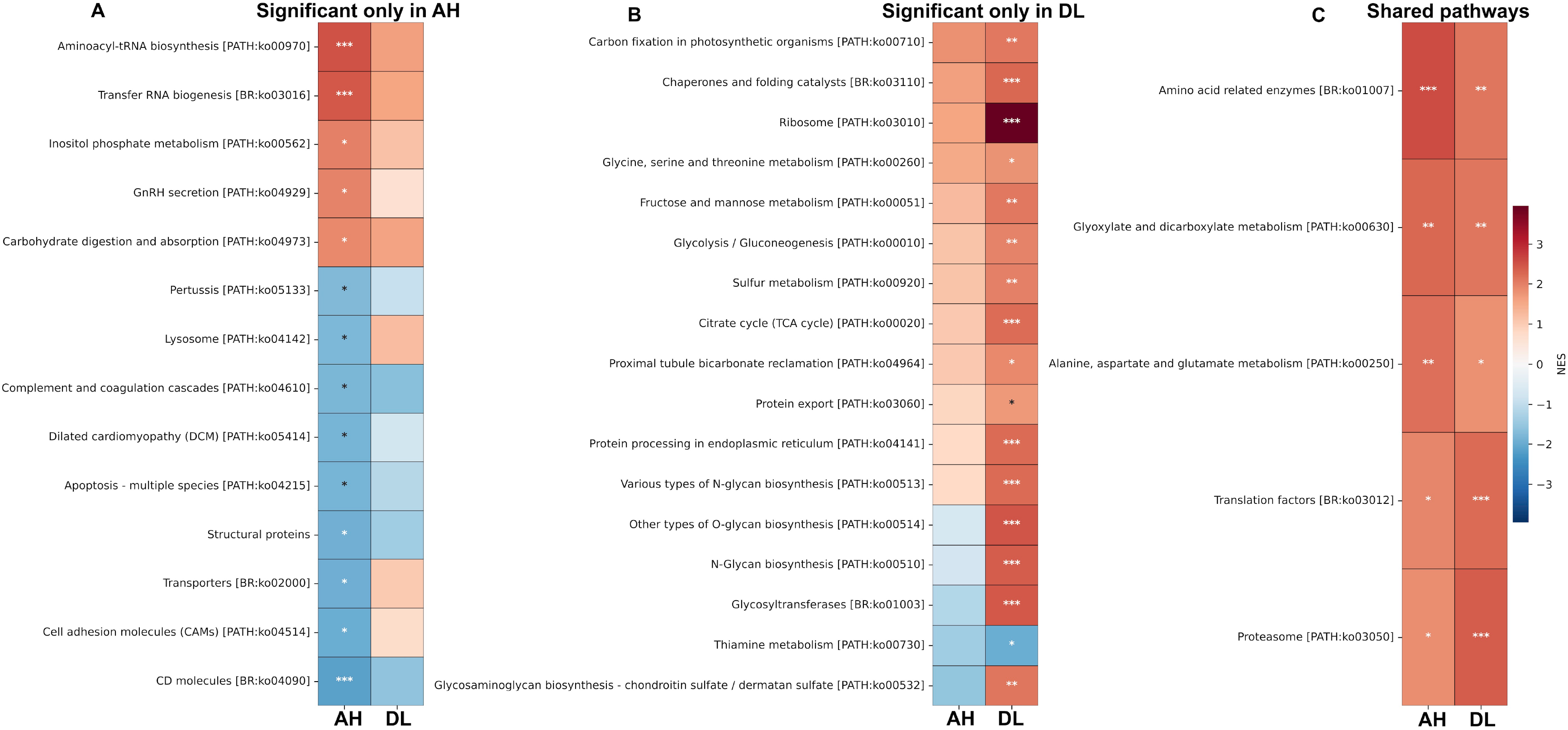
Functional pathway enrichment analysis of the proteome data. Heatmap shows significantly enriched KEGG pathways identified from utilizing Gene Set Enrichment Analysis. Pathways are separated into those significantly enriched: (**A)** only in AH, (**B)** only in DL, or **(C)** shared between both comparisons, when compared against HC. Colour intensity represents the normalized enrichment score (NES), with red indicating positive enrichment and blue indicating negative enrichment in the AH or DL treatments compared to HC. Asterisks denote the significance of pathway enrichment (* FDR *q-value <* 0.05, ** < 0.01 and *** < 0.001).

Whereas the apoptosis-multiple species pathway, which has been associated with SCTLD, was depleted (ko04215: NES = -1.83, *q*-value = 0.048). The lysosome pathway (ko04142; NES = -1.77, *q*-value = 0.050), as well as several immune system related pathways also depleted in AH relative to the HC samples, including, complement and coagulation cascades (ko04610; NES = -1.81, *q*-value = 0.042), CD molecules (ko04090; NES = -2.15, *q*-value < 0.001) and cell adhesion molecules (ko04515; NES = -2.16, *q*-value = 0.002).

A large shift was observed in DL relative to HC, with 22 pathways being enriched and only one depleted [**Figure 4B; Supplemental Table 4**]. Three pathways were associated with central carbon metabolism, including, the citrate cycle (TCA cycle) (ko00020; NES = 2.22, *q*-value < 0.001), fructose and mannose metabolism (ko00051; NES = 2.07, *q*-value = 0.003), and glycolysis/gluconeogenesis (ko00010; NES = 1.96, *q*-value = 0.007). We also observed the enrichment of five glycan synthesis pathways, which may be involved in coral wound healing and mucus production (Thobor et al., 2024). These included other types of O-glycan biosynthesis (ko00514; NES = 2.49, *q*-value < 0.001), glycosyltransferases (ko01003; NES = 2.44, *q*-value < 0.001), N-glycan biosynthesis (ko00510; NES = 2.39, *q*-value < 0.001), various types of N-glycan biosynthesis (ko00513; NES = 2.23, *q*-value < 0.001), and glycosaminoglycan biosynthesis-chondroitin sulfate/dermatan sulfate (ko00532; NES = 2.12, *q*-value = 0.002).

We also observed enrichment in five protein synthesis, folding, and transport pathways, suggesting a rapid proteomic response within the host. These included, the ribosome (ko03010; NES = 3.95, *q*-value < 0.001), chaperones and folding catalysts (ko03110; NES = 2.27, *q*-value < 0.001), protein processing in the endoplasmic reticulum (ko04141; NES = 2.24, *q*-value < 0.001), glycine, serine and threonine metabolism (ko00260; NES = 1.76, *q*-value = 0.033), and protein export (ko03060; NES = 1.70, *q*-value = 0.049). Lastly, we also observed enrichment in sulfur metabolism in DL samples relative to the healthy samples (ko00920; NES = 2.00, *q*-value = 0.005). Only the thiamine metabolism pathway was depleted in DL samples (ko00730; NES = -1.96, *q*-value = 0.042).

Five pathways associated with amino acid metabolism, protein synthesis, and protein turnover were enriched in both AH and DL tissues [**Figure 4C**]. These included amino acid related enzymes (ko01007; AH: NES = 2.58, *q*-value < 0.001; DL: NES = 2.12, *q*-value = 0.002), glyoxylate and dicarboxylate metabolism (ko00630; AH: NES = 2.27, *q*-value = 0.001; DL: NES = 2.10, *q*-value = 0.002), alanine, aspartate and glutamate metabolism (ko00250; AH: NES = 2.19, *q*-value = 0.002; DL: NES = 1.79, q-value = 0.026), translation factors (ko03012; AH: NES = 1.93, *q*-value = 0.024; DL: NES = 2.25, *q*-value < 0.001), and the proteasome (AH: NES = 1.83, *q*-value = 0.050; DL: NES = 2.46, *q*-value < 0.001).

## Discussion

### Overview of proteomic data

Samples were collected in triplicate from 10 coral colonies, comprising five healthy (HC) and five infected individuals that exhibited macroscopic SCTLD lesions. From each infected colony (IC), tissue was sampled both directly from the diseased lesion (DL) and from apparently healthy tissue (AH) located approximately 1-2 inches from the lesion. Proteomic data was generated for the coral host, algal endosymbionts, and the associated microbiome, with each component analyzed separately. Several factors limited usage of the endosymbiont and microbial datasets. First, healthy colonies contained variable endosymbiont communities dominated by *Durusdinium* and *Breviolum*, whereas infected colonies were dominated by *Symbiodinium necroappetens*, complicating direct comparisons between tissue types [**Figure 1**]. The presence of the opportunistic symbiont, *S. necroappetens* has been documented in corals undergoing stress and in dead tissue (LaJeunesse et al., 2015; Villela et al., 2025). In addition, endosymbiont cell counts were unavailable, preventing normalization of protein abundance values by symbiont cell abundance, an important consideration for comparative proteomics (Cunning & Baker, 2014; Nandi et al., 2026). In addition, the microbial proteome exhibited substantial missingness, with only three samples achieving > 50% protein completeness [**Supplemental Figure 2**]. Consequently, subsequent analyses focused exclusively on the coral host proteome, which exhibited substantially higher completeness (> 80%) across samples. Overall, after QC and filtering for outliers, 36 samples and 7,146 proteins remained for downstream analysis.

Within the host proteome, HC and AH samples were more closely associated in the PCA analysis than were DL samples, indicating that apparently healthy tissue from SCTLD affected colonies maintains a proteomic profile similar to that of healthy, unaffected colonies. In contrast, DL samples formed a distinct cluster, reflecting a conserved host proteomic response to infection [**Figure 2C, Supplemental Figure 1C**]. This pattern contrasts sharply with the microbial β-diversity values calculated from the same samples, in which the AH and DL tissues had more similar microbial community profiles compared to HC (Nandi et al., 2025). Together, these findings suggest that although the microbiome associated with apparently healthy tissue has already shifted toward a disease-associated state, the host proteome remains largely comparable to that of healthy colonies, potentially reflecting a delayed or buffered host response during early disease progression.

Differential abundance analysis supported this interpretation, with 480 DAPs identified between HC and DL, but only 12 DAPs between HC and AH, highlighting the extensive proteomic shifts associated with the active lesion, but not the disease-associated microbiome. At the pathway level, AH samples exhibited 10 enriched and 10 depleted pathways relative to HC, despite the small number of differentially abundant proteins. This likely reflects the greater sensitivity of GSEA, which enables the detection of coordinated, modest shifts across groups of functionally related proteins rather than relying solely on individual proteins exceeding significance thresholds (Joly et al., 2020). Together, these findings suggest that whereas apparently healthy tissue has not yet undergone widespread proteomic shifts, subtle, coordinated changes in biological processes are underway, potentially representing an early host response to disease before large-scale alterations in protein abundance become apparent in the data. These “early proteins” may prove useful for identifying markers of SCTLD and developing molecular diagnostic tools prior to full disease onset (Chille et al., 2025).

### Massive mobilization of the protein homeostasis machinery

We observed a pronounced enrichment of pathways associated with protein synthesis and homeostasis, in both AH and DL tissues [**Figure 4**]. Despite being visually indistinguishable from healthy tissues, AH tissues exhibited enrichment of aminoacyl-tRNA biosynthesis, tRNA biogenesis, translation factors, and proteasome pathways, suggesting that cells adjacent to lesions had already initiated an increased translational capacity and protein quality-control mechanisms [**Figure 4**]. This early response likely reflects the activation of the host in response to a pathogenic microbiome before visible lesion formation, consistent with the relatively small number of differentially abundant proteins detected in AH tissues. In contrast, these responses become substantially more pronounced in diseased tissues, where enrichment extended across the broader protein homeostasis network. In addition to the pathways enriched in AH, diseased tissues exhibited increased ribosomal activity, chaperones and folding catalysts, protein processing in the endoplasmic reticulum, and protein export, indicative of coordinated activation of protein synthesis, folding, trafficking, and degradation. Similar shifts in ribosomal and translational machinery have been reported in corals exposed to environmental stress suggesting that enhanced protein synthesis forms part of a broader cellular response to physiological stress (Mayfield et al., 2021; McRae et al., 2021; Oakley et al., 2017). These findings suggest that protein abundance remodelling was initiated before lesion formation and intensified during disease progression, likely to maintain cellular function under increasing physiological stress.

*Extracellular matrix maintenance appears to be a highly conserved response to SCTLD* Alterations to the extracellular matrix (ECM) have been implicated in multiple coral diseases (MacKnight et al., 2022; Traylor-Knowles et al., 2021), potentially affecting both tissue integrity and the coral surface microbiome layer, which plays an essential role in coral immunity (Glasl et al., 2016; Rosenberg et al., 2007; Shnit-Orland & Kushmaro, 2009).

Consistent with previous studies of SCTLD, DL tissues exhibited extensive remodelling of ECM-associated pathways and proteins following lesion formation (Rossin et al., 2026; Traylor-Knowles et al., 2021). At the pathway level, we observed enrichment of multiple glycan biosynthesis pathways, including N-glycan biosynthesis, O-glycan biosynthesis, and glycosaminoglycan biosynthesis. These pathways contribute to the synthesis and modification of glycoconjugates that comprise the extracellular matrix and mucins, which are the heavily glycosylated glycoproteins that comprise the coral surface mucus layer (Thobor et al., 2024). Increased activity of these pathways may therefore reflect enhanced production of mucus. We hypothesize that such a response could facilitate the shedding of pathogenic microbiome on the coral mucus, a proposed innate defence mechanism that limits microbial colonization and reduces pathogen load (Garren & Azam, 2012; Ritchie, 2006).

At the individual protein level, ECM-associated proteins exhibited a mixed response upon lesion formation [**Figure 3**]. Collagen-associated proteins displayed mixed patterns of abundance, whereas four hemicentin proteins were reduced. Previous SCTLD transcriptomic studies in *Orbicella faveolata* and *Montastraea cavernosa* reported extensive remodelling of ECM components but observed increased expression of laminin and fibronectin genes, which was interpreted as activation of wound-healing and tissue repair pathways (Papke et al., 2024; Traylor-Knowles et al., 2021). In contrast, we primarily observed reduced abundances of laminin, fibronectin, and several cytoskeleton-associated proteins in *D. labyrinthiformis*, suggesting that these reparative mechanisms may not be maintained in established lesions or may differ at the protein level compared to transcripts. Conversely, three copies of cartilage intermediate layer protein 1 increased in abundance, indicating selective maintenance of specific ECM components. These differences may reflect species-specific host responses to SCTLD, differences in disease stage, different polymicrobial etiologies, or the distinction between transcriptomic and proteomic responses. Species-specific responses are well documented in corals exposed to thermal stress, making similar variation in disease responses unsurprising (Da-Anoy et al., 2024; Molinari et al., 2025; Nandi et al., 2026). However, the overall loss of extracellular matrix integrity has been well documented in SCTLD lesions, consistent with our observation of reduced abundances of multiple ECM-associated proteins (Landsberg et al., 2020; Traylor-Knowles et al., 2021). The few proteins with increased abundance may likely be playing a role in wound healing which has been recorded in SCTLD responses (Papke et al., 2024).

### Known immune system responses appeared suppressed at the protein level

Corals mount diverse immune responses to pathogens through the activation of pattern recognition receptors, including lectins and integrins, followed by downstream effector responses such as complement activation, oxidative defences, and apoptosis (Palmer, 2018; Palmer & Traylor-Knowles, 2012; Papke et al., 2024; Parisi et al., 2020). Consistent with these observations, we found widespread changes in immune-associated pathways and proteins during SCTLD progression. In AH tissues, despite the presence of a pathogenic microbiome, pathway analysis revealed depletion of several immune-associated systems, including CD molecules, cell adhesion molecules, apoptosis, and complement and coagulation cascades [**Figure 4**]. These pathways were no longer significantly enriched or depleted in DL tissues following lesion formation.

With respect to protein abundance [**Figure 3]**, DL tissues exhibited pronounced changes in innate immune components. We observed decreased abundance of multiple pattern recognition proteins, including root-specific lectin-like, C-type lectin, intelectin, the mannose receptor CD206, and IFIH1, several of which have previously been implicated in coral disease responses, including SCTLD (Anderson et al., 2016; Fuess et al., 2018; Libro et al., 2013; Papke et al., 2024; Traylor-Knowles et al., 2021). In addition, immune effector proteins had decreased abundance, including two copies of complement C3 which has been previously associated with coral responses to bacterial infection (Brown et al., 2013; Van De Water et al., 2015; Wright et al., 2015). These observations suggest that innate immune defences are broadly diminished in lesion tissues of *D. labyrinthiformis*.

In addition to reductions in pathogen recognition and immune effector proteins, we observed widespread decreases in proteins associated with endosomal trafficking, and lysosomal transport, including DENND3, Rabaptin-5, RUFY1, FYCO1, TAX1BP1, SNF8, and LRSAM1. In corals, phagocytic cells engulf microorganisms and traffic them through phagosomal compartments for lysosomal degradation (Beavers et al., 2023; Helgoe et al., 2024.; Snyder et al., 2021). The coordinated decrease of proteins associated with these pathways may therefore reflect a reduced capacity for intracellular microbial clearance in DL tissues.

Lastly, previous studies have reported activation of apoptosis-associated pathways during SCTLD, suggesting that programmed cell death contributes to lesion progression and tissue remodeling (Beavers et al., 2023; McDonald, 2020; Papke et al., 2024; Traylor-Knowles et al., 2021). Pathway analysis indicated suppression of apoptosis in AH tissues. Furthermore, we did not observe widespread changes in apoptosis-associated proteins within established lesions. Instead, we observed decreased abundance of NOTCH1, a regulator of cell fate, proliferation, and apoptosis (Hemond et al., 2014; Marlow et al., 2012), together with decreased abundance of the apoptosis inhibitor baculoviral IAP repeat-containing protein (BIRC) (Kvitt et al., 2016). We also observed an increased abundance in the golgi associated plant pathogenesis protein, which acts as a negative regulator of autophagy (Daniels et al., 2015; Traylor-Knowles et al., 2021). Together, these observations support the hypothesis of dysregulation of apoptotic signaling rather than widespread activation of apoptosis.

Our findings raise an important question regarding the mechanisms underlying the apparent immune suppression. One possibility is that the response is host-mediated; i.e., corals suppress components of their innate immune system during the establishment and maintenance of symbiosis with Symbiodiniaceae (Mansfield & Gilmore, 2019; Valadez-Ingersoll et al., 2025). In this study, SCTLD-infected colonies were dominated by *S. necroappetens*, whereas healthy colonies contained more diverse mutualistic symbiont communities [**Figure 1C**]. It is therefore possible that colonization by *S. necroappetens* contributes to suppression of host immune pathways. An alternative, and not mutually exclusive, explanation is pathogen-mediated immune suppression. Numerous viruses can suppress host innate immunity by interfering with pathogen recognition and downstream immune signalling (Beachboard & Horner, 2016; Chan & Gack, 2016; Chathuranga et al., 2021). Given the increased abundance and apparent conservation of viral communities associated with SCTLD in multiple coral species (Nandi et al., 2025), viral modulation of host immunity represents a plausible hypothesis. However, the data presented in this study does not directly demonstrate a viral mechanism. Additional studies are required to determine whether pathogens actively contribute to immune suppression during SCTLD progression, or if they are simply colonizing an already immune-compromised coral.

### Differences between transcriptomic and proteomic results

Our findings highlight an important distinction between transcriptomic and proteomic responses under SCTLD. Previous transcriptomic studies reported activation of immune and apoptotic pathways following disease exposure, whereas our proteomic analysis of established lesions predominantly identified reduction in proteins involved in pathogen recognition, immune effector function, intracellular pathogen clearance, and regulation of apoptosis (Beavers et al., 2023, 2025; Papke et al., 2024; Traylor-Knowles et al., 2021). These differences likely reflect the complex relationship between transcription and protein abundance, which can be influenced by post-transcriptional regulation, protein turnover, pathogen-derived effects, the timing of sample collection, and coral species-specific responses (Chille et al., 2024; Da-Anoy et al., 2024; Molinari et al., 2025; Nandi et al., 2026). Inconsistency between observed changes in the transcriptome and proteome have been extensively demonstrated in corals and other systems (Gygi et al., 1999; Williams et al., 2023). These observations suggest that integrating transcriptomic, proteomic, and histological approaches is essential for resolving the temporal progression of SCTLD and disentangling early immune activation from the sustained dysfunction observed in advanced disease.

### Proteomic evidence points to host-endosymbiont dysbiosis

We observed the enrichment of central metabolic pathways in both AH and DL tissues. Notably, AH tissues exhibited enrichment of the carbohydrate digestion and absorption pathway, which may reflect a shift by the coral host toward greater heterotrophic nutrition in response to potentially reduced productivity of its endosymbionts (Grottoli et al., 2006). In DL tissues, we observed enrichment of several additional central metabolic pathways, including glycolysis, the TCA cycle, and fructose and mannose metabolism. Both tissue types showed enrichment of amino acid metabolism, including alanine, aspartate and glutamate metabolism, glycine, serine and threonine metabolism, as well as other amino acid-related enzymes. Similar metabolic shifts have been reported in corals exposed to thermal stress and bleaching, whereby increased amino acid metabolism accompanies enhanced protein synthesis and turnover (Montalbetti et al., 2026; Nandi et al., 2026; Williams et al., 2021). This observation is consistent with the enrichment of the ribosomal machinery observed in our dataset, suggesting increased protein production and metabolic remodelling during disease.

We also observed decreased abundances of proteins associated with the symbiosome and, consequently, coral-algal symbiosis. Specifically, we detected differential regulation of the lysosomal cathepsins, with decreased abundance of CTSL and increased abundance of CTSB, both of which have previously been implicated in coral stress responses, and symbiosome maintenance (Louis et al., 2017; Maruyama et al., 2026). The abundance of carbonic anhydrase proteins, which are essential for supplying carbon to the symbiosome and sustaining the photosynthetic activity of the endosymbionts was also reduced (Bertucci et al., 2013; Zoccola et al., 2015). In addition, several other lysosomal peptidases showed decreased abundance. Conversely, we observed increased abundances of GNMT and BHMT, proteins associated with coral stress and involved in one-carbon metabolism and methyl-group homeostasis (Aguilar et al., 2019; Nunn et al., 2025). We observed decreased abundance of HSP70, alongside increased abundance of the small heat shock protein HSPB6, suggesting a cellular stress response (Franzellitti et al., 2018; Rosic et al., 2011). Finally, proteins involved in reactive oxygen species (ROS) detoxification, including GST and GPX, were increased in abundance (Dias et al., 2019). These enzymes have been widely implicated in coral-algal dysbiosis and have also been proposed to contribute to immune defense through the mitigation of oxidative damage (Papke et al., 2024). These findings support the existing model that coral algal dysbiosis is a major component of SCTLD infection, although it remains to be elucidated if this is causative or correlative.

### Integrating host and microbiome responses during SCTLD progression

Based on host metaproteome and metagenome data (Nandi et al., 2025) determined from the same *D. labyrinthiformis* coral samples, a coherent model of SCTLD progression is emerging **[Figure 5]**. The metagenomic data demonstrate that both apparently healthy and diseased lesion tissue from infected colonies house dysbiotic microbial communities, whereas the present study reveals that these microbial shifts are accompanied by distinct host physiological responses. We observe changes in the algal endosymbiont community between infected and healthy colonies, suggesting that disruption of the coral-algal symbiosis is an early feature of disease progression.

**Figure 5:**
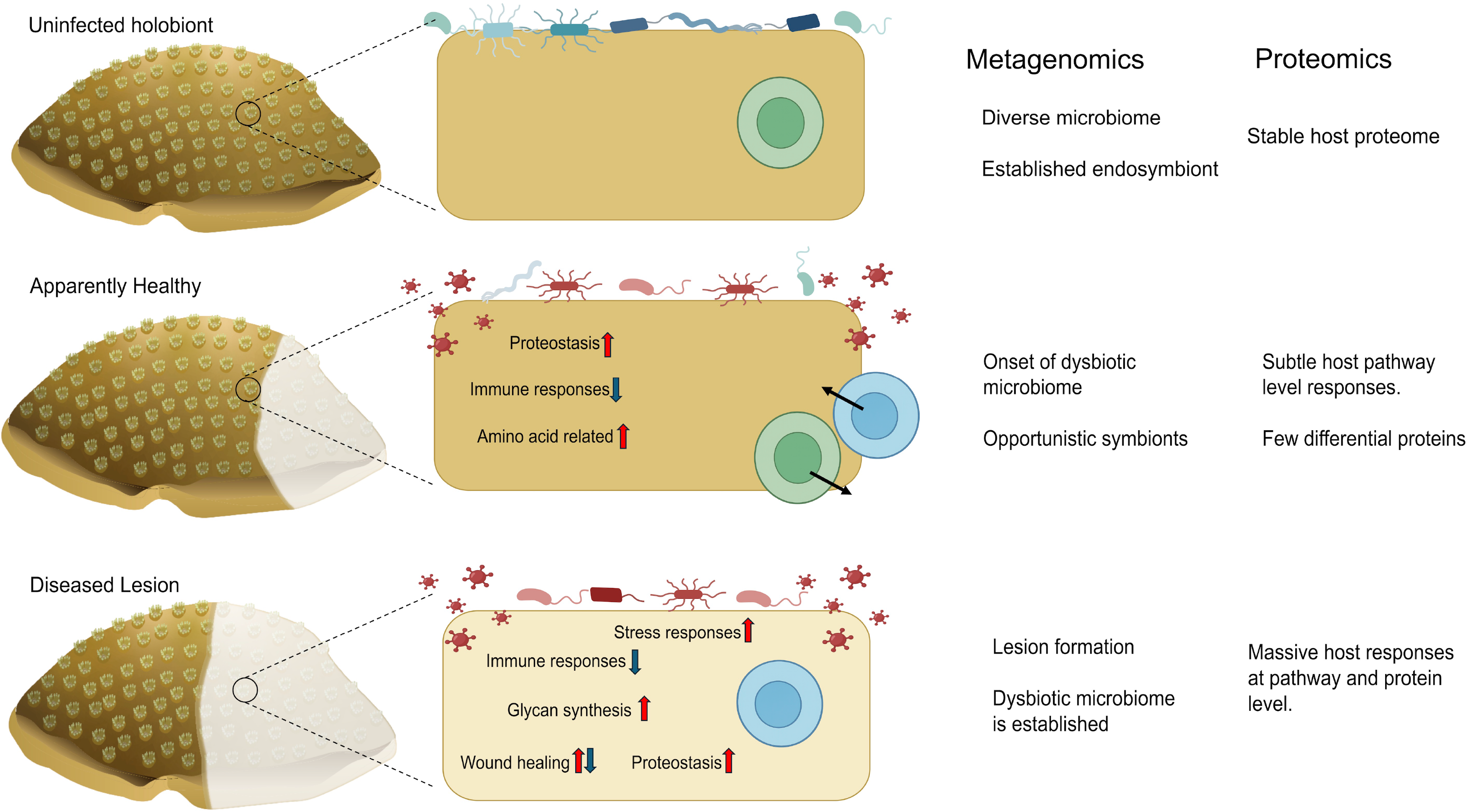
Multi-omics data-based model of SCTLD progression in *D. labyrinthiformis*. This is a testable hypothesis that integrates metagenomic and proteomic analyses in healthy, apparently healthy, and diseased lesion coral tissues. In the uninfected colony, a diverse microbiome coexists with a stable endosymbiont community, reflected by a stable host proteome. During the apparently healthy stage, the microbiome and virome shift toward dysbiosis with the emergence of opportunistic algal symbionts. In this stage, the host exhibits relatively subtle proteomic changes characterized by altered homeostasis, suppression of immune-associated pathways, and changes in amino acid metabolism. As the disease progresses to the lesion stage, a dysbiotic microbiome is established and extensive host proteomic remodelling occurs, including enhanced stress responses, altered glycan synthesis, disrupted wound healing and extracellular matrix, continued suppression of immune processes, and an increased homeostasis-related response. Our findings suggest that microbiome restructuring precedes widespread host proteomic reprogramming during SCTLD progression. Image prepared by Erin E. Chille.

In AH tissue, despite the presence of a dysbiotic microbiome, the coral host exhibits a broadly suppressed immune response together with enrichment of amino acid metabolism and heterotrophic metabolic pathways. These observations suggest that the host initiates a subtle yet clearly distinguishable proteomic response before the onset of visible lesions. One possible explanation is that SCTLD induces early coral-algal dysbiosis (Beavers et al., 2023; Rossin et al., 2026), either through direct impacts on the algal symbionts (Beavers et al., 2023; Work et al., 2021) or indirectly through pathogen-mediated disruption of the host. As dysfunctional symbionts are replaced (Cunning et al., 2015; Karrick et al., 2026), suppression of host immune pathways facilitates the establishment of new (potentially opportunistic) symbiotic partnerships (Mansfield et al., 2017; Mansfield & Gilmore, 2019; Valadez-Ingersoll et al., 2025). Alternatively, the observed immune suppression could reflect manipulation of host cellular processes by viruses, particularly given that immune modulation and host-pathway hijacking are well documented in many host-viral systems (Beachboard & Horner, 2016; Chan & Gack, 2016; Chathuranga et al., 2021). This latter, intriguing idea warrants further study.

As disease progresses to DL tissue, the host remains immunosuppressed but undergoes a marked shift towards a generalized stress response. We observe an increased abundance of glycan biosynthesis pathways, which may reflect enhanced mucus and mucin production (Thobor et al., 2024) as the coral attempts to physically remove or compartmentalize the dysbiotic microbial community. Concurrent increases in molecular chaperones, reactive oxygen species detoxification proteins, and proteins associated with cellular stress, together with widespread disruption of symbiosome-associated functions, suggest that the coral-algal symbiosis has largely collapsed by this stage (Beavers et al., 2023; Rossin et al., 2026).

We propose a model based on multi-omics data whereby SCTLD is characterized by an early phase of microbial dysbiosis and immune suppression that precedes visible tissue loss. This is followed by progressive breakdown of the coral-algal symbiosis and activation of generalized cellular stress responses during lesion development. Integrating host proteomics with microbiome and viral community dynamics provides a systems-level view of SCTLD progression and identifies several testable hypotheses to identify mechanisms of disease progression. These include early immune modulation, symbiosis destabilization, and potential viral involvement in SCTLD outbreak.

## Study Limitations and Future Directions

The proteomic workflow we utilized was optimized to target the coral host response, which thereby limited the recovery of microbial proteins. For future metaproteomic holobiont studies, we suggest that the microbial fraction be enriched prior to protein extraction sample-specific spectral libraries be used to reduce the large search space associated with metagenome-informed library-free DIA analysis. In addition, Symbiodiniaceae cell counts should be done to allow protein abundance normalization and additional physiological data such as photosynthetic efficiency should be gathered. Lastly, extending proteomic analysis to different SCTLD affected coral species will distinguish conserved disease responses from species-specific patterns, thereby aiding in the development of a molecular toolkit for disease diagnostics.

## Acknowledgements

Primary funding for this study was provided by the National Philanthropic Trust (23-7825575) in a grant awarded to Debashish Bhattacharya (DB). This work was also supported by grants from the National Science Foundation (2128073) and the USDA National Institute of Food and Agriculture Hatch Formula (NJ01180) awarded to DB. Samples were collected and exported to the US under permit number DJ-CON-1-2025-0005. We wish to acknowledge the captains of the research vessels who led the different sampling trips. This project is part of the International Climate Initiative (IKI). The Federal Ministry for the Environment, Nature Conservation and Nuclear Safety (BMU) supports this initiative based on a decision adopted by the German Bundestag.

## Data Accessibility and Benefit-sharing

All data are available in the main text or the supporting materials. The SCTLD mass spectrometry proteomics data have been deposited to the ProteomeXchange Consortium via the PRIDE partner repository with the dataset identifier XXXXXXX (https://www.ebi.ac.uk/pride/).

## Author Contributions

Debashish Bhattacharya, Shrinivas Nandi, Timothy G. Stephens, Maria F. Villalpando, Aldo Croquer (Conceptualization), Shrinivas Nandi, Timothy G. Stephens, Rebecca Garcia-Camps, Maria F. Villalpando, Aldo Croquer Haiyan Zheng (Investigation), Shrinivas Nandi (Visualization), Debashish Bhattacharya (Funding acquisition), Debashish Bhattacharya, Maria F. Villalpando, Rita I. Sellares-Blasco, Aldo Croquer (Supervision), Shrinivas Nandi (Writing – original draft), Debashish Bhattacharya, Shrinivas Nandi, Timothy G. Stephens, Kasey H. Walsh, Rebecca Garcia-Camps, Maria F. Villalpando, Rita I. Sellares-Blasco, Ainhoa L. Zubillaga, Haiyan Zheng, Aldo Croquer (Writing – review & editing).

## Supplemental Information

**Supplemental Figure 1: Summary of host proteomic data before cleaning. (A)** Bar chart representing the completeness of the host proteome in percent (x-axis) for the 44 samples (y-axis). (**B)** Shows the clustered heatmap of the proteome and samples prior to any cleanup during the *protii* clean up pipeline. Colored bars at the top of the heatmap shows the tissue type for the respective sample. (**C)** Principal component plot prior to any QC showing the sample distribution, highlighting outliers within the proteome.

**Supplemental Figure 2: Summary of microbiome protein completeness.** Bar chart representing the completeness of the microbiome proteome in percent (x-axis) for the 44 samples (y-axis).

**Supplemental Table 1:** Cleaned host proteomic data with log_2_ normalized abundance presented for the samples, along with differential abundance statistics generated using *protti*.

**Supplemental Table 2:** Differentially abundant proteins with statistics, KEGG term annotation and blast nr annotation.

**Supplemental Table 3:** Gene set enrichment analysis results for apparently healthy samples against healthy samples.

**Supplemental Table 4:** Gene set enrichment analysis results for diseased lesion samples against healthy samples.

